# Pairwise and Higher-Order Epistatic Interactions Have a Significant Impact on Bronchodilator Drug Response in African American Youth with Asthma

**DOI:** 10.1101/2020.03.04.977066

**Authors:** J Magaña, MG Contreras, KL Keys, O Risse-Adams, PC Goddard, AM Zeiger, ACY Mak, JR Elhawary, LA Samedy-Bates, E Lee, N Thakur, D Hu, C Eng, S Salazar, S Huntsman, T Hu, EG Burchard, MJ White

## Abstract

**Background:** Asthma is one of the leading chronic illnesses among children in the United States. Asthma prevalence is higher among African Americans (11.2%) compared to European Americans (7.7%). Bronchodilator medications are part of the first-line therapy, and the rescue medication, for acute asthma symptoms. Bronchodilator drug response (BDR) varies substantially among different racial/ethnic groups. Asthma prevalence in African Americans is only 3.5% higher than that of European Americans, however, asthma mortality among African Americans is four times that of European Americans; variation in BDR may play an important role in explaining this health disparity. To improve our understanding of disparate health outcomes in complex phenotypes such as BDR, it is important to consider interactions between environmental and biological variables.

**Results:** We evaluated the impact of pairwise and three-variable interactions between environmental, social, and biological variables on BDR in 617 African American youth with asthma using Visualization of Statistical Epistasis Networks (ViSEN). ViSEN is a non-parametric entropy-based approach able to identify interaction effects. We performed analyses in the full dataset and in sex-stratified subsets. Analysis in the full dataset identified six significant interactions associated with BDR, the strongest of which was an interaction between prenatal smoke exposure, age, and global African ancestry (IG: 1.09%, p=0.005). Sex-stratified analyses yielded additional significant, but divergent, results for females and males, indicating the presence of sex-specific effects.

**Conclusions:** Our study identified novel interaction effects significantly influencing BDR in African American children with asthma. Notably, we found that the impact of higher-order interactions was greater than that of pairwise or main effects on BDR highlighting the complexity of the network of genetic and environmental factors impacting this phenotype. Several associations uncovered by ViSEN would not have been detected using regression-based methods emphasizing the importance of employing statistical methods optimized to detect both linear and non-linear interaction effects when studying complex phenotypes such as BDR. The information gained in this study increases our understanding and appreciation of the complex nature of the interactions between environmental and health-related factors that influence BDR and will be invaluable to biomedical researchers designing future studies.

## Background

Asthma is an inflammatory disease of the lower respiratory tract, characterized by symptomatic difficulty of breathing in affected individuals.^1^ In the United States (U.S.), asthma is one of the leading chronic illnesses among children.^2^ Asthma is also the most disparate common disease in pediatric populations, with asthma prevalence, morbidity, and mortality rates varying widely by racial/ethnic group.^3^ Specifically, rates of asthma prevalence and mortality are two and four times higher, respectively, in African American children compared to European American children.^3^ Measures of asthma morbidity, including emergency department visits and missed school days, are also higher in African American children compared to their European American counterparts.^4^ Despite the higher asthma burden in the African American community, this population has been historically underrepresented in asthma research.^5,6^ Recent years have shown an increase in the inclusion of African Americans in large scale biomedical studies; however, this population is still comparatively understudied when contrasted with efforts aimed at European American populations.^5,6^

The disparity in asthma health outcomes across racial/ethnic groups may be due in part to a difference in drug response. Bronchodilators, specifically short acting β_2_-agonist medications such as albuterol, are the most commonly prescribed asthma medication in the United States. ^7,8^ Bronchodilator drug response (BDR) is the amount of airway obstruction that is reversible after the administration of bronchodilator medication. BDR varies significantly between racial/ethnic groups.^9-11^ Alarmingly, compared to other racial/ethnic groups, African American children with moderate-to-severe asthma respond poorly to bronchodilators, ranking second worst among all demographic groups.^11^

The estimated genetic heritability of bronchodilator drug response is approximately 28.5%.^12^ However, this estimate only represents the additive effect of each genetic factor on BDR variability and does not account for gene-environment, gene-gene, or variant-variant effects.

For example, while measures of air pollution, socioeconomic status, genetic ancestry, and obesity have all been independently associated with BDR and/or other asthma-related phenotypes, the amount of variation in BDR independently explained by each of these variables is relatively small, leaving a large portion of the variation in BDR undefined.^7,13-16^ A portion of the undefined variation in BDR is likely explained by gene-gene or gene-environment interactions.^17^ While these interactions could be linear or non-linear in nature, historically research in complex phenotypes, such as BDR, has traditionally focused primarily on the identification of linear interaction effects through the use of regression-based methods. Synergistic non-linear interactions (epistatic interactions) have recently been recognized as a significant source of variation underlying complex diseases.^18-20^ The widely employed regression-based models characteristic of large-scale genetic/epidemiological studies of complex disease, while adept at detecting linear, or additive, interaction effects, are not well powered to detect non-linear interactions. In addition, the majority of studies investigating interaction effects in complex diseases have utilized linear models in largely European populations.^21-23^ Consequently, there is a significant lack of research investigating epistatic interaction effects between environmental, psychosocial, demographic, and clinical factors in asthma research. Furthermore, the lack of genetic research in non-European populations perpetuates asthma health disparities, especially for those populations that carry a high disease burden, such as African Americans.

Visualization of Statistical Epistasis Networks (ViSEN) is a statistical program that optimizes detection of epistatic interactions through information-theoretic quantities, and is able to include multiple types of data as discrete random variables.^20,24^ ViSEN is also able to identify and quantify both linear and non-linear interactions and provide intuitive visualization of the potentially complex relationships between large numbers of variables using a network-based approach. Additionally, ViSEN has been shown to be more powerful than standard regression-based methods in detecting interaction effects, suggesting that ViSEN is a promising investigative tool that can be applied to studies of complex diseases such as BDR.^20,24-28^

In this study, we conducted pairwise (two-variable) and higher order (three-variable) interaction analyses using ViSEN to study the impact of both linear and epistatic interactions between social, psychological, and biological variables on BDR in 617 African American youth with asthma. We then performed these analyses in sex-stratified subsets to identify sex-specific effects in our study population. Our study is the first to rigorously interrogate clinical, environmental, and demographic information to identify hidden non-linear interactions affecting BDR in African American children with asthma. By identifying previously overlooked epistatic interactions that significantly influence BDR in African American youth with asthma; our study aims to provide novel information that can aid in characterizing targets for future health intervention strategies and improving the design of future studies of BDR.

## Results

### Study Participants

The SAGE study included African American participants aged 8 to 21 years with and without asthma. Individuals without asthma and individuals missing any phenotype data were excluded from our analysis (Table 1), yielding a total of 617 individuals with asthma (Males = 335, Females = 282). In the full dataset, ViSEN analysis identified two variants independently associated with BDR status: sex (p = 0.01) and perceived experience of discrimination (*p* = 0.03) (Table 1). All marginal effects of single variables, defined by ViSEN as main effects, identified by ViSEN were also detected using standard descriptive statistics (Table 1). Similarly, we explored the possibility of main effects in sex-stratified subsets of our population. For males, no independent effects were identified, while in females, global African ancestry was significantly associated with BDR responder status (p = 0.02, Additional File 1, Additional File 2).

**Table 1.** Study Demographics.

|  |  |  |  | ViSEN | Descriptive Statistics |
| --- | --- | --- | --- | --- | --- |
| Categorical Variable |  | BDR Responders | BDR Non-Responders | p-value<sup>#</sup> | p-value |
| Sample Size, N |  | 171 | 446 | --- | --- |
| Sex (% Female) |  | 37% | 49% | **0.01** | **0.01** * |
| Age, yrs. (Mean, [SE]) |  | (14, [0.275]) | (14, [0.169]) | 0.93 | 0.77<sup>+</sup> |
| Body Mass Index | Obese | 63 | 142 | 0.26 | 0.28* |
|  | Non-Obese | 108 | 304 |  |  |
| Experience of Discrimination | Yes | 97 | 208 | **0.04** | **0.03** * |
|  | No | 74 | 238 |  |  |
| Prenatal Smoke Exposure | Yes | 33 | 86 | 1.00 | 1.00* |
|  | No | 138 | 360 |  |  |
| Socioeconomic Status | > Low | 116 | 290 | 0.57 | 0.57* |
|  | Low | 55 | 156 |  |  |
| Air Pollution (NO<sub>2</sub>) | ≥ Median | 91 | 217 | 0.35 | 0.36* |
|  | < Median | 80 | 229 |  |  |
| Global African Ancestry | ≥ 80% | 116 | 269 | 0.11 | 0.10* |
|  | < 80% | 55 | 177 |  |  |

Summary statistics for all phenotypic data included for analysis in this study are presented above. Significant p-values are highlighted in **bold**. *p-values* represent the significance of the independent effects, or main effects, of specified variables on BDR responder status. <sup>#</sup>p-value calculated from ViSEN's Mutual Information (MI) Test. MI is a metric that quantifies the reduction in uncertainty about the distribution of one variable given an understanding of the other; \* p-value calculated from Chi-squared Test of Independence; <sup>+</sup> p-value calculated from linear regression

### ViSEN Pairwise Interaction Effects

We identified five significant pairwise (two-variable) interactions associated with BDR in the full dataset using ViSEN. ViSEN calculates the strength of an interaction’s impact on BDR using an information-theory metric known as Information Gain (IG) (see Methods Section). Pairwise interaction models significantly associated with BDR, in order of strength defined by Information Gain (IG) were: [1] prenatal smoke exposure and socioeconomic status (SES) (IG = 0.91%, p = 0.007), [2] experience of discrimination and SES (IG = 0.54%, p = 0.025), [3] age and body mass index (IG = 0.54%, p = 0.028), [4] sex and global African ancestry (IG = 0.49%, p = 0.036), and [5] experience of discrimination and prenatal smoke exposure (IG = 0.46%, p = 0.045) (Table 1A and Figure 1). Of these five significant pairwise interaction models, two models ([1] prenatal smoke exposure and SES (β = −3.484, p = 0.007) and [2] age and body mass index (β = 2.577, p = 0.014)) were also identified using linear regression analysis (Table 1A).

**Figure 1.**
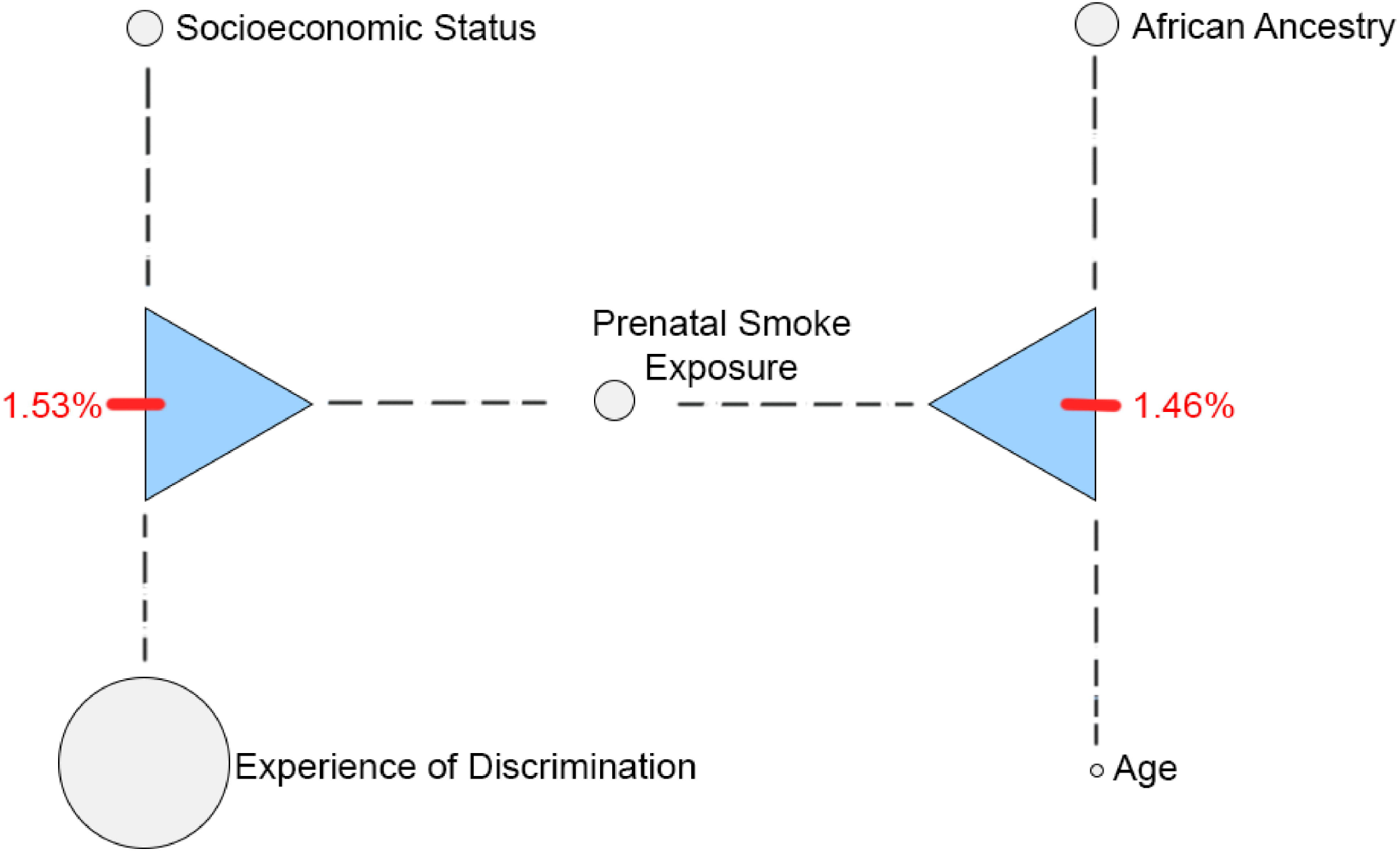
Visualization of Interaction Network in the Full Dataset. Visual representation of the interaction network for all significant interaction effects identified by ViSEN for the full dataset. The size of an individual node corresponds to the amount of Mutual Information (MI) resulting from the independent main effects of each variable. The strength of significant pairwise interactions corresponds to the thickness of the lines connecting independent nodes along the network. Triangles and dotted lines represent a significant three-way interaction effect between variables.

ViSEN analysis of pairwise interactions performed in sex-stratified subsets of our study population revealed disparate significant pairwise interaction effects among females and males. In females, the three strongest pairwise interaction effects included: [1] prenatal smoke exposure and SES (IG = 3.62%, p < 0.001), [2] experience of discrimination and SES (IG = 2.19%, p = 0.008), and [3] experience of discrimination and global African ancestry (IG = 2.05%, p = 0.008) (Table 2B, Figure 2). Of the three significant pairwise interaction effects uncovered by ViSEN, none were detectable using linear regression analysis (Table 2B). There were no significant pairwise interaction models identified by either ViSEN or linear regression in the male-only subset analysis.

**Table 2.** ViSEN Interaction Models Significantly Associated with BDR in the Full Dataset.

**A. Full Dataset**
| Variable 1 | Variable 2 | ViSEN |  | Linear Regression |  |
| --- | --- | --- | --- | --- | --- |
| | | IG | p-value | $\beta$ | p-value |
| Prenatal Smoke Exposure | SES<sup>+</sup> | 0.91% | 0.007 | -3.484 | **0.007** |
| Experience of Discrimination | SES<sup>+</sup> | 0.54% | 0.025 | -0.656 | 0.532 |
| Age | BMI<sup>*</sup> | 0.54% | 0.028 | 2.577 | **0.014** |
| Sex | African Ancestry | 0.49% | 0.036 | -0.148 | 0.886 |
| Experience of Discrimination | Prenatal Smoke Exposure | 0.46% | 0.045 | 1.012 | 0.423 |

**B. Sex-Stratified Subsets**
| Subset | Variable 1 | Variable 2 | ViSEN |  | Linear Regression |  |
| --- | --- | --- | --- | --- | --- | --- |
| | | | IG | p-value | $\beta$ | p-value |
| FEMALE | Prenatal Smoke Exposure | SES<sup>+</sup> | 3.62% | < 0.001 | -3.118 | 0.110 |
|  | Experience of Discrimination | SES<sup>+</sup> | 2.19% | 0.008 | -1.469 | 0.327 |
|  | Experience of Discrimination | African Ancestry | 2.05% | 0.008 | 0.628 | 0.680 |
| MALE | --- | --- | --- | --- | --- | --- |

Information Gain (IG) and $p$ -value results for significant interaction models identified by ViSEN with corresponding linear regression results. Significant linear regression $p$ -values are highlighted in **bold**. (A) ViSEN results for significant pairwise interactions in the full dataset. (B) ViSEN results for significant pairwise interaction in sex-stratified subsets of the full dataset. All ViSEN and linear regression models were adjusted for age, baseline lung function (Pre-FEV<sub>1</sub>), and marginal effects of each independent variable included in the specified interaction model; analyses performed in the full dataset were also adjusted for sex. BMI<sup>\*</sup>: Body Mass Index; SES<sup>+</sup>: Socioeconomic Status; $\beta$ : Beta value from linear regression model

**Figure 2.**
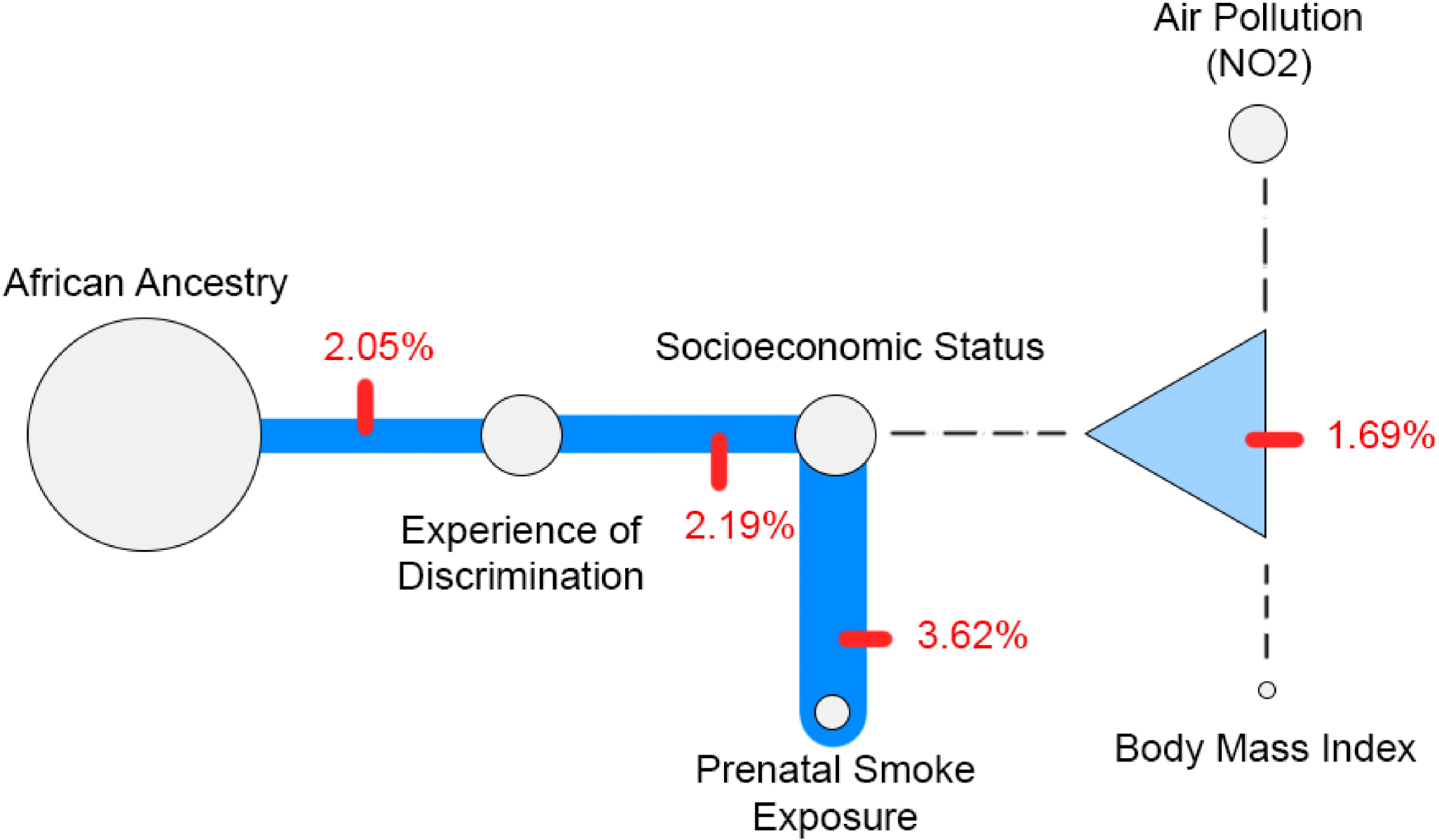
Visualization of Interaction Network in Females. Visual representation of the interaction network for all significant effects generated in ViSEN for female only subset. The size of an individual node corresponds to the amount of Mutual Information (MI) resulting from the independent main effects of each variable. The strength of significant pairwise interactions corresponds to the thickness of the lines connecting independent nodes along the network. Triangles and dotted lines represent a significant three-way interaction effect between variables.

### ViSEN Higher-Order (Three-Variable) Interaction Effects

We identified four significant higher-order (three-variable) interactions associated with BDR in the full dataset using ViSEN. Higher-order interaction models significantly associated with BDR, in order of strength defined by IG, were: [1] prenatal smoke exposure, age, and African ancestry (IG = 1.09%, p = 0.005), [2] experience of discrimination, sex, and African ancestry (IG = 0.85%, p = 0.011), [3] prenatal smoke exposure, sex, and SES (IG = 0.74%, p = 0.023), and [4] prenatal smoke exposure, age, and NO_2_ air pollution (IG = 0.65%, p = 0.047) (Table 3A and Figure 1). Of these four significantly associated interaction models, only one model (prenatal smoke exposure, age, and African ancestry) was also identified using linear regression analysis (β= 6.268, p= 0.021) (Table 3A).

**Table 3.** Higher-Order Interaction Models Significantly Associated with BDR identified by ViSEN.

**A. Full Dataset**
| Variable 1 | Variable 2 | Variable 3 | ViSEN |  | Linear Regression |  |
| --- | --- | --- | --- | --- | --- | --- |
| | | | IG | p-value | $\beta$ | p-value |
| Prenatal Smoke Exposure | Age | African Ancestry | 1.09% | 0.005 | 6.268 | **0.021** |
| Experience of Discrimination | Sex | African Ancestry | 0.85% | 0.011 | -1.386 | 0.501 |
| Prenatal Smoke Exposure | Sex | SES<sup>+</sup> | 0.74% | 0.023 | -0.442 | 0.867 |
| Prenatal Smoke Exposure | Age | Air Pollution (NO<sub>2</sub>) | 0.65% | 0.047 | 5.487 | 0.089 |

**B. Sex-Stratified Subsets**
| Dataset | Variable 1 | Variable 2 | Variable 3 | ViSEN |  | Linear Regression |  |
| --- | --- | --- | --- | --- | --- | --- | --- |
| | | | | IG | p-value | $\beta$ | p-value |
| FEMALE | Air Pollution (NO<sub>2</sub>) | BMI<sup>*</sup> | SES<sup>+</sup> | 1.69% | 0.026 | -6.791 | **0.032** |
| MALE | Prenatal Smoke Exposure | Experience of Discrimination | SES<sup>+</sup> | 1.53% | 0.018 | 2.818 | 0.436 |
|  | Prenatal Smoke Exposure | Age | African Ancestry | 1.46% | 0.024 | 7.756 | **0.035** |

Information Gain (IG) and $p$ -value results for significant interaction models associated with BDR identified by ViSEN with corresponding linear regression results. Significant linear regression $p$ -values are highlighted in **bold**. All ViSEN and linear regression models were both adjusted for age, sex, baseline lung function (Pre-FEV<sub>1</sub>), and marginal effects (what ViSEN calls “main effects”) of each variable included in the specified interaction model. (A) ViSEN results for three-way interaction in the full dataset. (B) ViSEN results for significant three-way interactions in sex-stratified subsets of the full dataset. BMI<sup>\*</sup>: Body Mass Index; SES<sup>+</sup>: Socioeconomic Status; $\beta$ : Beta value from linear regression model

ViSEN analysis of three-variable interactions performed in sex-stratified subsets of our study population generated discordant results. In females, a single significant three-variable interaction model containing NO_2_ air pollution, body mass index, and SES (IG = 1.69%, p = 0.026; β= −6.791, p = 0.032) was detected by both ViSEN and linear regression analysis (Table 3B, Figure 2). Conversely, there were two significant three-variable interaction models identified by ViSEN in the male-only subset analysis: [1] prenatal smoke exposure, experience of discrimination, and SES (IG = 1.53%, p=0.018) and [2] prenatal smoke exposure, age, and African ancestry (IG = 1.46%, p=0.024) (Table 3B, Figure 3). The interaction model in males containing prenatal smoke exposure, age, and African ancestry was also found to be significantly associated with BDR using linear regression analysis (β= 7.756, p= 0.035) (Table 3B).

**Figure 3.**
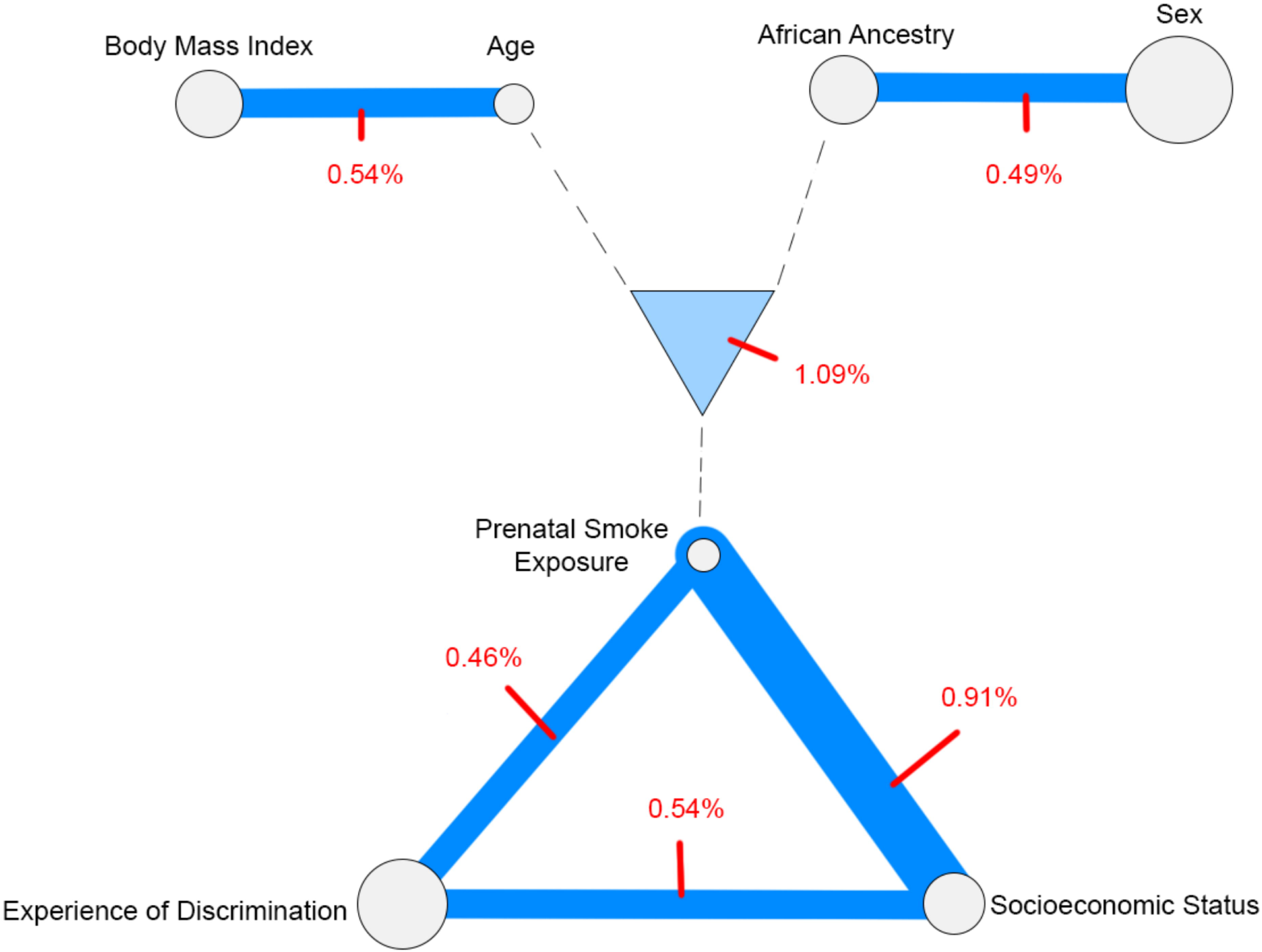
Visualization of Interaction Network in Males. Visual representation of the interaction network for all significant effects generated in ViSEN for the male-only subset. The size of an individual node corresponds to the amount of Mutual Information (MI) resulting from the independent main effects of each variable. Triangles and dotted lines represent a significant three-way interaction effect between variables.

## Discussion

The high inter-individual variability of BDR between racial/ethnic populations may contribute to disparities in asthma morbidity and mortality observed in African American children with asthma. An existing pharmacogenomic study has characterized genetic components which may explain response to albuterol in this population.^29^ Related studies have also noted ethnic-specific and other phenotypic differences in bronchodilator drug responsiveness.^9,11,30^ However, few have observed the joint effect of variables influencing BDR. To the best of our knowledge, our study is the first to analyze the interaction between clinical, genetic, environmental, and psychosocial factors affecting drug response in African American children and adolescents using a non-parametric method optimized to detect epistatic interactions. Here we demonstrate an integrative, investigative approach, which identifies novel gene-environment epistatic interactions that may influence variability of BDR.

ViSEN consistently identified novel pairwise and higher-order interactions occurring within our study population that, to our knowledge, have not been discussed elsewhere. Notably, the most informative interaction occurring in females between prenatal smoke exposure and socioeconomic status (IG = 3.62%) was discovered completely independent of main effects for either variable. This was also true in our full dataset where the interaction between prenatal smoke exposure and socioeconomic status (IG = 0.91%) was also identified in the absence of main effects for either variable. Importantly, of the nine significant interaction effects identified by ViSEN in the full dataset, only three effects of these effects were detected using linear regression. This trend was also seen in our sex stratified subset analyses; linear regression detected only one of the four significantly associated interactions identified in females, and only one of the two significantly associated interactions models uncovered by ViSEN in males. This suggests that the majority of the interaction effects revealed by ViSEN are synergistic (non-linear) in nature and therefore hidden when assessed using standard regression-based methods. It is also worth noting that the interactions not detected by linear regression typically displayed greater Information Gain (IG) metrics compared with those that were found by linear regression, further implying that non-linear interactions may have a stronger impact on BDR than linear models.

Another interesting result of our study was the discordance of results between male and female only subsets. Our results suggest the presence of significant sex-specific differences in the interactions affecting BDR status. Another important revelation of our study was the strong effect of higher-order interactions on BDR. In the full dataset, a three-variable interaction model had the strongest and most significant association with BDR (Table 3A). This suggests that incorporating the study of more complex models may lead to novel discoveries in this complex phenotype.

Interpreting results from ViSEN entails several considerations. The first consideration involves the nature of the variables and their method of collection. BDR represents a clinical continuous measurement obtained via spirometry, but other variables such as an experience of discrimination were collected via a self-reported questionnaire. Therefore, there is a possibility that measurement error contributed to the detection of interactions within this study, especially in variables that are not easily validated by repeated measurements. However, in our study we have rigorously identified classifications for each variable and validated our measurements by either consulting clinical guidelines or referencing previous literature.^13,14,31-34^ All phenotype data included in this study has also been successfully used in other studies relating to asthma phenotypes.^13,14,31-34^ Again, it is paramount to consider that regardless of the rigor of data collection, ViSEN must collapse continuous variables into a ranked form and therefore some information will be lost as result.

It should also be noted that while ViSEN can identify linear and non-linear interactions and quantify the amount of information provided by these models, it does not provide the directionality of these effects (i.e. whether interactions contribute positively/negatively to BDR responsiveness). To mitigate this limitation for studies in which information on directionality of identified interaction effects would be useful, it may be possible to supplement ViSEN analyses with additional post-hoc analyses such as quantitative multifactor dimensionality reduction (qMDR) or the more familiar Dunn test, to aid in further characterization of identified interaction effects.^35,36^ However, it should be noted that while supplementation of ViSEN analysis in this way is possible, results from these post-hoc tests may not be easily interpretable for every phenotype or interaction model. It will be the responsibility of individual researchers to determine if their specific study lends itself to this type of further analyses.

## Conclusion

Investigating epistatic gene-environment interactions is important to understanding variation in BDR in the context of asthma outcomes, such as morbidity and mortality, and improving health equity across the U.S. Social determinants of health are recognized as some the most predictive factors contributing to individual health outcomes. Therefore, the inclusion of variables describing our “built environment” (i.e. psychosocial factors such as socioeconomic status, experiences of discrimination, etc.) in gene-environment interaction studies of BDR is crucial. Our study incorporated biological, environmental, and psychosocial factors into a single comprehensive analysis of pairwise and higher-order interaction models impacting BDR in African American youth with asthma. We identified novel interaction models significantly impacting BDR in a population that carries a high disease burden (increased asthma morbidity and mortality compared to European American children with asthma) and has been historically understudied and underserved.

Parametric methods, such as generalized linear modeling, are the statistical tool of choice in most scientific efforts to characterize interactions associated with BDR and other complex biological phenotypes. The strength of using parametric methods is that they are well-understood and therefore generally interpretable. However, methodology that is powered to detect only additive linear relationships makes assumptions about the normality of variable sample distributions and the distributional relationship of any identified interactions that may not hold true for many complex traits. We contend that a significant portion of gene-environment interactions effecting complex phenotypes like BDR are non-additive in nature, and the results of this study support this theory. The diversity and complexity of the interactions impacting BDR should be embraced by employing non-parametric methods such as ViSEN that are optimized to identify multiple types (linear/non-linear) gene-environment interactions that might go undetected by less inclusive methods.

The nonparametric nature of ViSEN facilitates analyses that are more inclusive of the varying relationships and interactions between health factors likely impacting complex clinical phenotypes versus traditional linear modeling as we highlight in this study of BDR. Our study represents a collaboration of computer science, biology, environmental epidemiology, and social epidemiology that may lead to increased knowledge of asthma drug response and asthma morbidity in an “at-risk” population. We believe that further collaboration between these fields is necessary to fully understand the mechanism of action and impact of each interaction so that we incorporate novel findings into actionable medical intervention that benefits all patients.

## Methods

### Study Population

The Study of African Americans, Asthma, Genes, & Environments (SAGE) is a case-control study consisting of 1,710 participants ranging from ages 8 to 21 years old recruited from the San Francisco Bay Area between 2008 and 2014. The SAGE study protocol and patient population have been previously described in further detail, elsewhere.^14,29,31^ Briefly, all SAGE participants included in this study self-identified as African American, as did their parents and all four grandparents. All participants presented no history of other lung or chronic non-allergic illnesses upon study enrollment. Trained interviewers administered questionnaires to the participants and/or the parents/caretakers of the participants to collect basic demographic information, medical histories, and environmental exposure-related information.^14^

Our study consisted of 617 participants with physician-diagnosed asthma from SAGE, with complete data on sex, age, global African genetic ancestry, body mass index (BMI), any experience of discrimination, socioeconomic status, prenatal smoke exposure, ambient NO_2_ exposure over the first year of life, and BDR (Additional File 3). Our sex-stratified subsets consisted of 335 and 282 male and female individuals, respectively (Additional File 1 and Additional File 2). Appropriate descriptive statistics for participants in the full dataset, as well as sex-stratified subsets, were generated using the R statistical computing environment (Table 1, Additional File 1, and Additional File 2). ViSEN does not accommodate continuous variables. Consequently, the outcome variable and all explanatory variables included in interaction analyses were dichotomized prior to analysis as described below and in Additional File 3.

### Bronchodilator Drug Response (BDR)

The primary outcome of this study is bronchodilator responder status. Responder status was determined from individual spirometry measurements taken before and after administration of albuterol. Following American Thoracic Society recommendations, pulmonary function was measured prior to albuterol administration and then repeated 15 minutes after administration of four puffs (90 μg/puff) of albuterol.^37^ This process was repeated a third time after a second dosage of albuterol: two puffs for participants under the age of 16 and four puffs for older participants.^29^ Asthma medications were withheld from participants 12 hours before spirometry.^38^ BDR (ΔFEV_1_) was calculated as the mean percentage change in measured Forced Expiratory Volume (FEV_1_) before and after albuterol administration, using the post-albuterol spirometry with the maximal change ((post-FEV_1_ – pre-FEV_1_) / pre-FEV_1_) x 100%. For each participant in this study, BDR (ΔFEV_1_) was used to classify bronchodilator responder status as either a responder, ≥ 12%, or a non-responder, < 12%.^34^ We excluded two participants who were statistical outliers for BDR (raw values) as previously described.^39^

Using a previously published protocol, deviance residual values generated from a generalized logistic model (BDR∼ age + sex +pre-FEV_1_) were employed to adjust dichotomized BDR values by age, sex, and lung function.^28^ Deviance residual values for BDR resulting from the GLM were then used to reclassify individual responder status in the following manner: residual values ≥ 0% were reclassified as responders; residual values < 0% were reclassified as non-responders. Ultimately, reclassified responder status was utilized as the outcome of interest for ViSEN analysis. Individual BDR status was fully concordant before and after covariate adjustment

### Age

The participant’s age was calculated as the difference between the age of enrollment (the date on the eligibility form) and the date of birth. Discrete variables ranked from 0-1 were generated depending on whether an individual was aged below (ranked 0), or at and above the median for the population (ranked 1).

### Global African Ancestry

Participants included in this study were previously genotyped using the Axiom^®^ LAT1 array (World Array 4, Affymetrix, Santa Clara, CA).^40,41^ For every individual, we estimated the genetic proportion contributed by an African ancestral population. These estimates, obtained using an unsupervised run of ADMIXTURE, were considered as an average over each individual’s entire genome to comprise a global ancestry variable.^42^ Reference haplotypes of African and European individuals used in ADMIXTURE were gathered from the HapMap phase III YRI and CEU populations.^15^ In this study, values for global African ancestry were dichotomized according to their distribution above or equal to (1) or below (0) the U.S. national average of 80% global African ancestry.^43^

### Body Mass Index

At the time of enrollment, each study participant was measured via a calibrated scale and stadiometer for weight (kg) and height (m), respectively. Body mass index percentile values were subsequently calculated through the following formula: BMI = (kg)/ (m^2^). BMI percentile values were generated using guidelines for BMI categories from the U.S. Centers for Disease and Control and Prevention Growth Charts. BMI percentile values were dichotomized as either 0 or 1 depending on whether they fell below (< 95%) or above/equal to (≥ 95%) the Obese BMI classification.^16^

### Perceived Experience of Discrimination

Self-reported racial/ethnic discrimination was ascertained using the Experiences of Discrimination Questionnaire.^44^ Consistent with previous studies, we included questions pertaining to our population: “Have you ever experienced discrimination, been prevented from doing something, or been hassled or made to feel inferior, in any of the following situations because of your race, ethnicity, color, or language? (1) At School; (2) Getting medical care; (3) Getting services in a store or restaurant; and (4) On the street or in a public setting”; with choice for each question of *Yes* or *No*.^45,46^ Experiences of discrimination were dichotomized as none or any (affirmative answer to at least one situation). Interviewers required permission of caretakers to administer questions to participants equal to or less than 16 years of age. Perceived experiences of discrimination were reported at time of recruitment.^46^

### Prenatal Smoking

Prenatal exposure to smoke was determined from questionnaire information regarding the self-reported smoking status of participant’s mother during pregnancy. Binary values were assigned for smoking status based on whether the mother was a non-smoker (0) or active smoker (1) during the pregnancy of the participant.

### Socioeconomic Status

We created a composite index for socioeconomic status (SES) derived from three socioeconomic indicators: mother’s educational attainment, insurance status, and household income as previously described.^32^ Each component variable was independently assigned a value scored on a three-point scale ranging from low income (0), to medium income (1), to high income (2). Finally, for the purpose of our study, individuals were classified as either having a low (0) or medium/high (1) composite socioeconomic scores.

### Nitrogen Dioxide Exposure

TomTom/Tele Atlas EZ-Locate software (TomTom, Amsterdam, The Netherlands) was utilized to assign geographic coordinates for each participant’s residential history. We collected regional ambient air pollution data from the US Environmental Protection Agency Air Quality System based on these geographic coordinates.^13^ Measures of average ambient NO_2_ exposure (µg/ppb) were estimated over the first year of each participant’s life. If the participant moved during this period, NO_2_ exposure was weighted depending on the number of months spent at each residence. Discrete binary variables with values 0 or 1 were generated depending on whether the individual was exposed to below (0) or greater than/equal to the median NO_2_ exposure (1) for the sample population within the first year of life.

### Visualization of Statistical Epistasis Networks (ViSEN)

ViSEN is a statistical program used to perform network-based analyses that quantify and visualize pairwise and higher-order epistatic interactions. Research suggests that ViSEN is more powerful than standard regression-based methods for detecting nonlinear, non-additive interaction effects.^19,26,33^ Effects of single explanatory variables, which ViSEN defines as main effects, on phenotype status is calculated using Mutual Information (MI). MI, derived from information theory, quantifies the reduction in uncertainty about the distribution of one variable given an understanding of the other. ViSEN measures the strength of interaction effects on phenotype in terms of Information Gain (IG); IG is an information-theoretical metric that quantifies how much additional phenotypic variance is explained by jointly considering two or more variables versus an additive model of their individual effects. We calculated pairwise (two-variable) and higher order (three-variable) interaction effects on BDR in the full dataset and in gender-stratified subsets. To assess the significance of IG values computed from pairwise and higher order interactions in our dataset, permutation (n=1000) was performed. Permutation datasets were created by randomly shuffling BDR responder status. For each permuted dataset, the IG was recomputed for every pairwise and higher order interaction model to form a null distribution of IG values. *p-values* for IG calculations were generated by comparing the number of permutation-based IG values equal to or larger than the IG value observed in our real dataset. In our study, we considered interaction models with permutation p-values < 0.05 as statistically significant.

### Assessment of ViSEN identified Interactions using Linear Regression

ViSEN has been shown to be more powerful than standard methods in identifying epistatic interactions. However, since most previous interaction studies employed common statistical methods to identify associations, it is possible that non-linear interactions that significantly impact phenotype were overlooked.^20^ Following the standard assumption in biomedical studies of additive main effects and multiplicative interaction effects, we created multiplicative interaction terms to reflect each significant interaction model identified by ViSEN. We then investigated whether any of the newly created interaction terms was significantly associated with BDR using linear regression. For linear regression analysis BDR was left as a continuous variable, and regression models were adjusted for age, sex, and baseline lung function (pre-FEV_1_), consistent with ViSEN models. Also consistent with ViSEN models, regression models were additionally adjusted for the independent effects of all variants included in the interaction terms as well as lower order interaction models included in the interaction term if applicable. Linear regression models assessing pairwise interactions were coded as follows: BDR ∼ age + sex + FEV1 + variable1 + variable2 + variable1*variable2 (pairwise interaction term). Linear regression models assessing higher-order (three-variable) interaction terms were coded as follows: BDR ∼ age + sex + FEV1 + variable1 + variable2 + variable3 + variable1*variable2 + variable2*variable3 + variable 1*variable3 + variable1*variable2*variable3 (three-variable interaction term). All regression analyses were performed in R.^47^

## Abbreviations

BDR: Bronchodilator drug response
MI: Mutual Information
IG: Information Gain
ViSEN: Visualization of Statistical Epistasis Networks

## Declarations

### Ethics approval and Consent to Participate

A total of 617 participants from the SAGE study were used to generate the results described in this manuscript. The SAGE study was approved by the Institutional Review Board of the University of California San Francisco (Laurel Heights Panel) (IRB# 10-02877; Reference # 271317). All participants 18 years of age at the time of study enrollment provided their written consent to participate in this study. Parents of participants under 18 years of age provided their written assent for the participation of their children in this study.

### Consent for publication

Not Applicable

### Availability of data and materials

Biological, environmental and phenotypic data analyzed in the current study are available in the dbGAP repository (study accession number phs00921.v1.p1). Psychosocial data analyzed in the current study (experience of discrimination and socioeconomic status) are not publicly available due to the sensitive nature of the data and privacy concerns for study participants. Psychosocial data is currently stored in the UCSF Box repository and is available from the corresponding author upon reasonable request (https://ucsf.box.com/s/2cx8v52u1ouql02w8io3mhwe2tzell5m).

### Authors’ Contributions

J.M. and M.J.W. designed the current study, drafted the manuscript, and analyzed and interpreted the data. K.L.K. and M.G.C. made substantial contributions to data interpretation and drafted the manuscript. A.C.Y.M., D.H., N.T., C.E., S.H., J.R.E., and S.S. generated covariate data and substantively revised the manuscript. O.R.A., P.G.C., A.Z., L.A.S.B., and E.L. made substantial contributions to manuscript revisions. T.H. interpreted the data, computed ViSEN p-values, and made substantial revisions to the manuscript draft. E.G.B. conceived and designed the SAGE study cohort, facilitated the acquisition of clinical and environmental data analyzed in the manuscript, and made substantial revisions to the manuscript draft.

## Acknowledgements

The authors wish to acknowledge the following SAGE co-investigators for subject recruitment, sample processing and quality control: Luisa N. Borrell, DDS, PhD, Emerita Brigino-Buenaventura, MD, Adam Davis, MA, MPH, Michael A. LeNoir, MD, Kelley Meade, MD, Fred Lurmann, MS and Harold J. Farber, MD, MSPH. The authors also wish to thank the staff and participants who contributed to the SAGE study.

## Additional Files

File name: Additional File 1

File format: .docx

Title of data: Supplemental Table 1. Female Subset Demographics

Description of data: Demographic information for female-only subset analyses

File name: Additional File 2

File format: .docx

Title of data: Supplemental Table 2. Male Subset Demographics

Description of data: Demographic information for male-only subset analyses

File name: Additional File 3

File format: .docx

Title of data: Supplemental Table 3. Phenotypic data included for analysis in this study

Description of data: Description and categorization of data included in ViSEN analyses

