## Additional File 3 for "Pairwise and Higher-Order Epistatic Interactions Have a Significant Impact on Bronchodilator Drug Response in African American Youth with Asthma"

**Supplemental Table 3.** Phenotypic data included for analysis in this study.

| *Variable Name* | *Classification* | *Ranking* |
| --- | --- | --- |
| Age | < sample median | 0 |
|  | ≥ sample median | 1 |
| Air Pollution (NO_2_) | < sample median | 0 |
|  | ≥ sample median | 1 |
| Body Mass Index | Non-Obese | 0 |
|  | Obese | 1 |
| Bronchodilator Drug Response (BDR) | BDR Non-Responder : < 12% ΔFEV_1_^*^ | 0 |
|  | BDR Responder: ≥ 12% ΔFEV_1_^*^ | 1 |
| Experience of Discrimination | No Experience of Discrimination | 0 |
|  | Any Experience of Discrimination | 1 |
| Global African Ancestry | < 80% Ancestry | 0 |
|  | ≥ 80% Ancestry | 1 |
| Prenatal Smoke Exposure | Mother Non-Smoker | 0 |
|  | Mother Active Smoker | 1 |
| Sex | Female | 0 |
|  | Male | 1 |
| Socioeconomic Status | Low | 0 |
|  | Medium or High | 1 |

^*^ΔFEV_1_ was calculated as described in Methods.
