## Additional File 2 for "Pairwise and Higher-Order Epistatic Interactions Have a Significant Impact on Bronchodilator Drug Response in African American Youth with Asthma"

|  | | | | **ViSEN** | **Descriptive Statistics** |
| --- | --- | --- | --- | --- | --- |
| Categorical Variable | | BDR  Responders | BDR  Non-Responders | p-value^#^ | p-value |
| Sample Size, N | | 107 | 228 | --- | --- |
| Age, yrs.  (Mean, [SE]) | | (13, [0.327]) | (14, [0.225]) |  | 0.51^+^ |
| Body Mass Index | Obese | 39 | 65 | 0.18 | 0.18^*^ |
|  | Non-Obese | 68 | 163 |  |  |
| Experience of Discrimination | Yes | 60 | 102 | 0.06 | 0.07^*^ |
|  | No | 47 | 126 |  |  |
| Prenatal Smoke Exposure | Yes | 18 | 44 | 0.75 | 0.69^*^ |
|  | No | 89 | 184 |  |  |
| Socioeconomic Status | > Low | 71 | 156 | 0.71 | 0.80^*^ |
|  | Low | 36 | 72 |  |  |
| Air Pollution (NO_2_), µg/ppb | ≥ Median | 53 | 101 | 0.43 | 0.44^*^ |
|  | < Median | 54 | 127 |  |  |
| Global African Ancestry | ≥ 80% | 66 | 135 | 0.67 | 0.76^*^ |
|  | < 80% | 41 | 93 |  |  |

**Supplemental Table 2. Male Subset Demographics**

Summary statistics for all phenotypic data included for analysis in this study are presented above. Significant p-values are highlighted in **bold**. *p-value*s represent the significance of the independent effects, or main effects, of specified variables on BDR responder status. ^#^p-value calculated from ViSEN’s Mutual Information (MI) Test. MI is a metric that quantifies the reduction in uncertainty about the distribution of one variable given an understanding of the other; ^*^ p-value calculated from Chi-squared Test of Independence; ^+^ p-value calculated from linear regression
