## Additional File 1 for "Pairwise and Higher-Order Epistatic Interactions Have a Significant Impact on Bronchodilator Drug Response in African American Youth with Asthma"

|  | | | | **ViSEN** | **Descriptive Statistics** |
| --- | --- | --- | --- | --- | --- |
| Categorical Variable | | BDR  Responders | BDR  Non-Responders | p-value^#^ | p-value |
| Sample Size, N | | 64 | 218 | --- | --- |
| Age, yrs.  (Mean, [SE]) | | (15, [0.477]) | (14, [0.250]) | 0.89 | 0.45^+^ |
| Body Mass Index | Obese | 24 | 77 | 0.74 | 0.86^*^ |
|  | Non-Obese | 40 | 141 |  |  |
| Experience of Discrimination | Yes | 37 | 106 | 0.22 | 0.25^*^ |
|  | No | 27 | 112 |  |  |
| Prenatal Smoke Exposure | Yes | 15 | 42 | 0.48 | 0.58^*^ |
|  | No | 49 | 176 |  |  |
| Socioeconomic Status | > Low | 45 | 134 | 0.21 | 0.25^*^ |
|  | Low | 19 | 84 |  |  |
| Air Pollution (NO_2_), µg/ppb | ≥ Median | 38 | 116 | 0.40 | 0.47^*^ |
|  | < Median | 26 | 102 |  |  |
| Global African Ancestry | ≥ 80% | 50 | 134 | **0.02** | **0.02*** |
|  | < 80% | 14 | 84 |  |  |

**Supplemental Table 1. Female Subset Demographics**

Summary statistics for all phenotypic data included for analysis in this study are presented above. Significant p-values are highlighted in **bold**. *p-value*s represent the significance of the independent effects, or main effects, of specified variables on BDR responder status. ^#^p-value calculated from ViSEN’s Mutual Information (MI) Test. MI is a metric that quantifies the reduction in uncertainty about the distribution of one variable given an understanding of the other; ^*^ p-value calculated from Chi-squared Test of Independence; ^+^ p-value calculated from linear regression
